# Recurrent Neural Network-based Prediction of O-GlcNAcylation Sites in Mammalian Proteins

**DOI:** 10.1101/2023.08.24.554563

**Authors:** Pedro Seber, Richard D. Braatz

**Affiliations:** Massachusetts Institute of Technology, 77 Massachusetts Avenue, Cambridge, MA 02139

## Abstract

O-GlcNAcylation has the potential to be an important target for therapeutics, but a motif or an algorithm to reliably predict O-GlcNAcylation sites is not available. In spite of the importance of O-GlcNAcylation, current predictive models are insufficient as they fail to generalize, and many are no longer available. This article constructs MLP and RNN models to predict the presence of O-GlcNAcylation sites based on protein sequences. Multiple different datasets are evaluated separately and assessed in terms of strengths and issues. The models trained in this work achieve considerably better metrics than previously published models, with at least a two-fold increase in F_1_ score relative to previously published models; the specific gains vary depending on the dataset. Within a given dataset, the results are robust to changes in cross-validation and test data as determined by nested validation. The best model achieves an F_1_ score of 36% (more than 3.5-fold greater than the previous best model) and a Matthews Correlation Coefficient of 35% (more than 4.5-fold greater than the previous best model), and, for the F_1_ score, 7.6-fold higher than when not using any model. Shapley values are used to interpret the model ‘s predictions and provide biological insight into O-GlcNAcylation.

## 1 Introduction

Glycosylation is a co- and post-translational modification in which a glycan or glycans are added to proteins. When a glycan is added to the oxygen of an amino acid (typically serine or threonine), this process is called O-linked glycosylation. When the first glycan added is an N-Acetylglucosamine (GlcNAc), this process is called O-GlcNAcylation [1]. O-GlcNAcylation is mediated by the enzymes OGT and OGA and is important functionally and structurally [1] [2]. Conversely, incorrect glycosylation or deglycosylation is associated with multiple diseases such as cancers [3], infections [4], and congenital disorders [5].

Glycosylation has great potential for diagnoses and treatments, which may benefit the biomed-ical and pharmaceutical industry, physicians, and patients. For example, many forms of carcinoma show increases in fucosylation, branching, and sialylation [6]. The vast majority of neuroblastomas express disialoganglioside; as such, neuroblastoma may be treated by anti-disialoganglioside mono-clonal antibodies according to Phase I–III studies [7] [8]. Antibodies containing poly-*α*2,8-sialylation have increased half-lives without the introduction of tolerance problems [9]. Conversely, therapeu-tics may be hindered by glycans not produced by humans. An example is the immunogenic N-glycolylneuraminic acid [10], which is in some CHO-cell-derived glycoproteins [11]. Recent research has shown O-GlcNAcylation can be a powerful target for therapeutics [12], further highlighting the potential of glycosylation.

In spite of how important glycosylation is for biotherapeutics, some challenges remain. Analyses need to take into account the locations of glycosylation sites and the glycan compositions at each of these sites [6]. It is challenging to investigate specific functions of glycosylation due to the wide diversity of glycan sites and distributions [1]. The complex equipment required to determine glycosylation prevents clinical laboratories from widely analyzing patient samples, restricting personalized medicine [6].

Computational tools can aid researchers in better understanding and predicting glycosylation patterns. Models can be classifiers, such as YinOYang (YoY) [13] or O-GlcNAcPRED-II [14], or regressors, such as the models in Refs. [15], [16], and [17]. In the context of glycosylation, classifiers may predict whether an amino acid can be N- or O-glycosylated. Regressors may quantitatively predict the glycan distribution of a glycosylation site. Models to predict the presence of O-GlcNAcylation sites have insufficient performance to be helpful tools. A 2021 review (Ref. [18]) found that no published model can achieve a precision *≥* 9% on a medium-sized independent dataset, indicating O-GlcNAcylation prediction models fail to generalize successfully despite the high metrics that they can achieve in their respective training datasets. These models also have low F_1_ scores and Matthew Correlation Coefficients (MCCs), further indicating their performance is lacking.

In this work, we construct recurrent neural network (RNN) classification models to predict the presence of O-GlcNAcylation sites from mammalian protein sequence data. The model construction procedures employ cross-validation and rigorous unbiased prediction error estimation. Our RNN models achieve significantly higher metrics than previously reported models, and their predictions are interpreted through Shapley values. Open-source software is provided so that other researchers can reproduce the work, retrain the models as additional or higher quality O-GlcNAcylation data become available, and use the models to further improve the understanding of O-GlcNAcylation.

## 2 Materials and Methods

### 2.1 Datasets

Three experimental datasets, one at a time, are used to construct the models. Table 1 summarizes the size and features of each dataset. The first dataset, named “Ref. [18] – Original” in this publication, is taken directly from Ref. [18]. This dataset contains human-selected descriptors based on sequence and structure, but does not contain the protein sequence directly, and has certain issues, such as repeated entries, which likely lead to test-set leakage. Thus, a second dataset, named “Ref. [18] – Modified” in this publication, is built from the raw data in Ref. [18]. This second dataset contains only protein sequences for site prediction. A third, larger dataset, named “Ref. [19] – Modified” in this publication, is built from the processed data from Ref. [19]. The dataset was modified to remove non-mammalian proteins, remove proteins without site information, and split entries with multiple isoforms. Due to the presence of isoforms and homologous proteins, care was taken to not include the same sequence multiple times in the processed dataset. This selection was done based on a window size of 5 AA on each side of the central S/T (11 AA total) even for the larger windows, reducing any effects due to site similarity. 20% of each dataset was separated for testing, with the remaining 80% used for cross-validation with five folds. Thus, test-set leakage is avoided. To ensure robustness against variations in the data and avoid biases due to our selection of training and testing sets, a 5-fold nested validation is performed with the best model in this work. In each round of nested validation, 20% of the data are separated as the round ‘s test set. The other 80% are used in five-fold cross-validation for hyperparameter selection (as above). The best model is evaluated against that round ‘s test set.

**Table 1:**
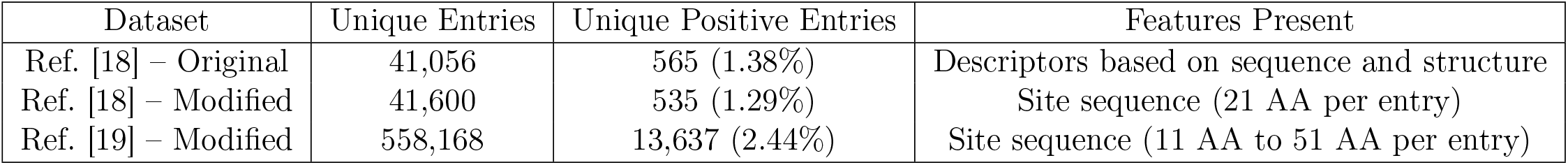
Summary of the properties of the datasets used in this work. Datasets came from Refs.[18] and [19] and are modified by us when noted.

### 2.2 Artificial Neural Networks (ANNs)

Multilayer perceptrons (MLPs) are constructed for the Ref. [18] – Original dataset, as it did not contain sequence information, while recurrent neural network (RNN) models are constructed for the other datasets. Model construction was done using PyTorch [20] and other Python packages [21] [22] [23]. For the MLP models, 32 different layer configurations, 4 different learning rates (10^*−*2^, 5*×*10^*−*3^, 10^*−*3^, 5*×*10^*−*4^), 3 different activation functions (ReLU, tanh, and tanhshrink), and varying loss weights for the positive class were used. For the RNN models, 2 LSTM size configurations (selected based on the number of MLP features in the original dataset), 7 different MLP layer configurations, 2 different learning rates (10^*−*2^, 5*×*10^*−*3^), 2 different activation functions (ReLU and tanhshrink), and varying loss function weights for the positive class were used.^1^ Moreover, the RNNs trained with the Ref. [19] dataset used an AdamW optimizer with a weight decay parameter of *λ* = 10^*−*2^ and cosine scheduling [24]. Cosine scheduling has been used primarily in the computer vision field and achieves great results in the context of imbalanced datasets [25] [26]. The best hyperparameters for the MLP or each RNN size are determined by a grid search, testing each combination of layers, learning rate, and activation function. The combination with the highest cross-validation average F_1_ score is selected and, for each RNN size, its performance is reported for an independent test dataset. To interpret the model’s predictions, Shapley values are calculated using the shap Python package [27] [28]. These interpretable values are also evaluated against the same test set.

### 2.3 Model Evaluation Metrics

Binary classification models emit two predictions. Because the real data have two potential categories, there are a total of four categories in which a prediction may fall: true positive (TP), false positive (FP), true negative (TN), and false negative (FN). There are multiple metrics that combine some or all of these categories to assess the quality of a model. The simplest of these metrics is accuracy (Eqn. S1), defined as the number of correct guesses divided by the total guesses. However, accuracy is not a suitable metric for imbalanced datasets, as it is possible to achieve high accuracy by simply always predicting the majority class. To correct this issue, two other simple metrics can be used. Recall (a.k.a. sensitivity, hit rate, or true positive rate) is the number of true positives divided by all positives in the real data (Eqn. S2). Precision (a.k.a. positive predictive value) is the number of true positives divided by all elements classified as positive by the model (Eqn. S3). Because recall and precision are opposed to each other, the F_1_ score metric was created to balance both and allow assessment of models with a single metric (Eqn. S4). The negative-class equivalents of recall and precision are the specificity (a.k.a. selectivity or true negative rate) and the negative predictive value; however, these metrics are not used in this work because it focuses on the discovery of positive O-GlcNAcylation sites. Moreover, the negative imbalance of the datasets makes these last two metrics less suitable for evaluation.

The most used metric to assess binary classification models in multiple fields is the area under the receiver operating characteristic curve (ROC-AUC) [29]. However, the ROC-AUC metric suffers from many issues and can lead to overoptimistic and incorrect assessments, especially when working with negatively-imbalanced data [29] [30] [31]. To analyze the quality of binary classification models over multiple thresholds, it is recommended to use the precision-recall (P-R) curve instead [29] [32][33] [34]. Finally, to capture in a single metric all four categories in which a prediction may fall, the Matthews correlation coefficient (MCC), also called the (Yule) phi coefficient, may be used (Eqn. S5). As a correlation coefficient, the MCC falls between -1 and 1 instead of the typical 0 and 1.

## 3 Results

### 3.1 MLP Models Considerably Surpass Previously Published Models in Terms of Precision, F_1_ Score, and MCC on the Original Dataset of Ref. [18]

All of the data-driven models constructed in this study for the prediction of O-GlcNAcylation sites are trained with hyperparameters selected by cross-validation. Using the “Ref. [18] – Original” dataset, this section compares our MLP model with previously published models: YinOYang [13], O-GlcNAcPRED-II [14], OGTSite [35], and the models in Ref. [18].

The central thesis of Ref. [18] is that O-GlcNAcylation prediction models fail because no model reviewed in that article achieved a precision greater than 9%. However, that conclusion was based on the low precision of the specific models evaluated in that reference. At low, less strict acceptance thresholds, our MLP behaves similarly to YinOYang (Fig. 1). Beginning from a threshold equal to 10^*−*6^ or higher, our MLP displays greater precision at the same recall level. The precision continues to increase monotonically with threshold, while the F_1_ score peaks at a model threshold equal to 0.8. At its maximum F_1_ score, our MLP model has a 151% improvement in the F_1_ score, a 307% in precision relative to the best former models (with a total precision of 35.3%), and a 90.9% improvement in MCC (Table 2). As such, our MLP model shows that the central thesis of Ref. [18] is not valid and that predictive models of the location of O-GlcNAcylation sites can be constructed with reasonable precision while also surpassing previous models in other metrics.

**Table 2:** Recall, precision, F_1_ score, and MCC metrics (in %) for the points with highest F_1_ score for different O-GlcNAcylation site prediction models. Model “YinOYang” is from Ref. [13], Model “O-GlcNAcPRED-II” is from Ref. [14], Model “OGTSite” is from Ref. [35], and Model “Mauri” is the best model from a collection of models from Ref. [18]. The metrics for these previously published models are from Tables 2 and 4 of Ref. [18]. Model “Our MLP” comes from this work. “No Model” sets all sites to potentially positive – that is, sets the threshold to 0.

| Metric | YinOYang<br>(From [13]) | O-GlcNAcPRED-II<br>(From [14]) | OGTSite<br>(From [35]) | Mauri<br>(From [18]) | Our MLP<br>(This Work) | No Model |
| --- | --- | --- | --- | --- | --- | --- |
| Recall (%) | 14.36 | 65.09 | 49.09 | 39.82 | 22.64 | 100 |
| Precision (%) | 6.89 | 3.90 | 6.20 | 3.10 | 35.29 | 1.38 |
| $F_1$ Score (%) | 9.31 | 7.36 | 11.01 | 5.75 | 27.59 | 2.72 |
| MCC (%) | 8.10 | 11.73 | 14.44 | 7.03 | 27.56 | 0.00 |

**Figure 1:**
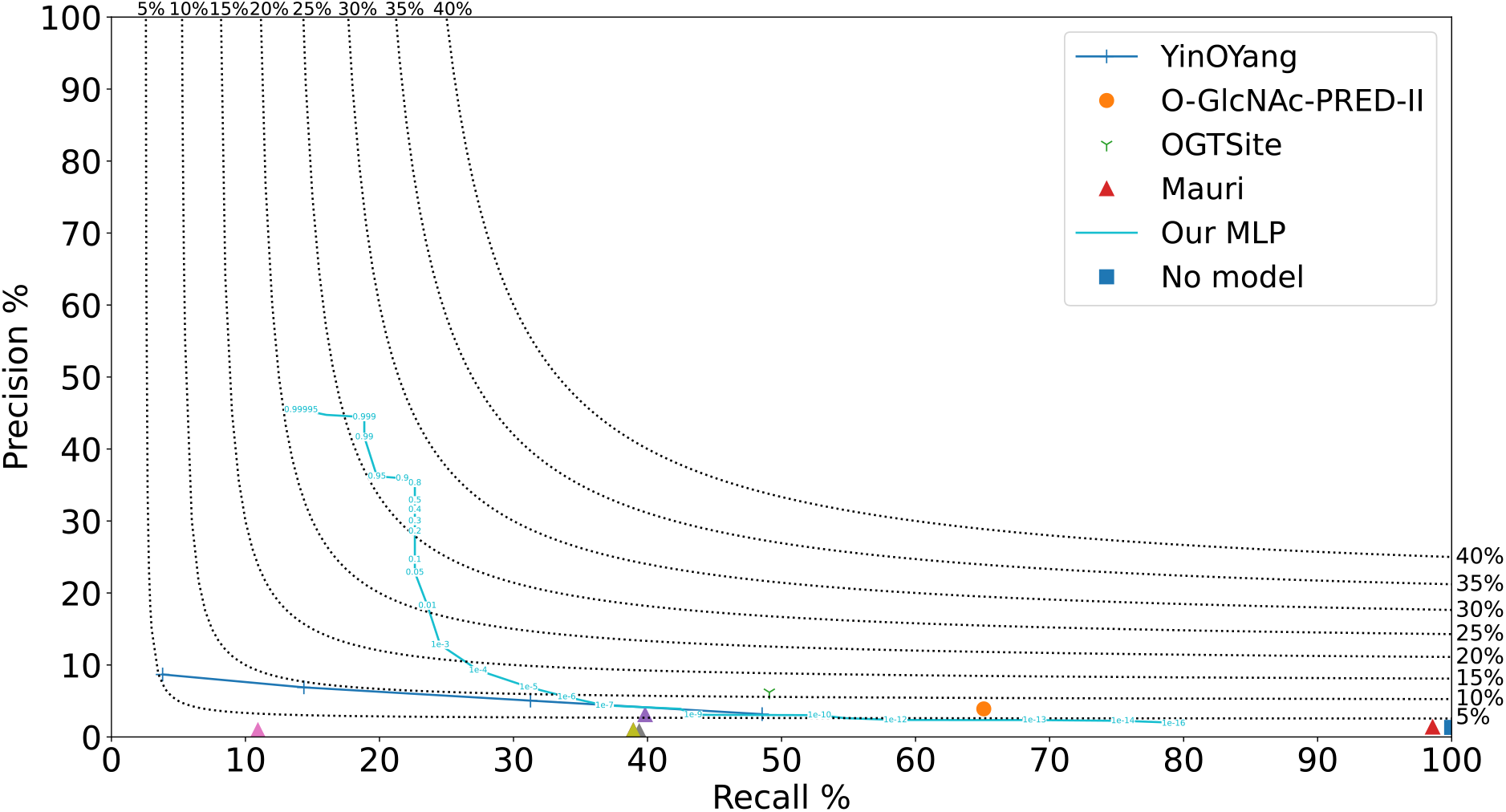
P-R curves for O-GlcNAcylation site prediction models tested on the original dataset of Ref. [18]. The black dotted lines are isolines of the F_1_ score, as labeled on the top and right sides of the plot. Model “YinOYang” is from Ref. [13], Model “O-GlcNAcPRED-II” is from Ref. [14], and Model “OGTSite” is from Ref. [35] (complete P-R curves are not shown because the models are no longer publicly available). Models “Mauri” are a set of models from Ref. [18]. Metrics for these previously published models are from Tables 2 and 4 of Ref. [18]. Model “Our MLP” comes from this work; the numbers on its curve represent the minimum threshold for a site to be considered positive. “No Model” sets all sites to potentially positive – that is, sets the threshold to 0.

After the MLP was fully trained, we noticed that the “Ref. [18] – Original” dataset has some issues, such as entries that did not match the raw data and repeated entries, which may lead to test-set leakage and overoptimistic predictions. Moreover, the models made predictions based on human-selected descriptors, which may be incomplete or biased and are not trivial to obtain, making model usage inconvenient for the end-user. To remedy these issues, we constructed a new dataset from the raw data of Ref. [18], which is called “Ref. [18] – Modified” in this work. This modified dataset uses sequence data instead of human-selected descriptors.

On that corrected dataset, our RNNs behave similarly to YinOYang at low thresholds (Fig. S1). Beginning from a threshold of 10^*−*10^ for the RNN-76 model and 10^*−*7^ for the RNN-152 model^2^, our RNNs display greater precision at the same recall level. The precision for both RNN models continues to increase nearly monotonically with increasing threshold, while the F_1_ score peaks at a threshold of 0.99 for RNN-76 and 0.999 for RNN-152. Moreover, the RNN-76 is strictly superior to the RNN-152 for all thresholds *≥* 10^*−*10^. At its F_1_ score maximum, the RNN-76 model has a 134% improvement in the F_1_ score, a 391% improvement in precision, and a 136% improvement in MCC relative to YinOYang ^3^ (Table S1).

The performance of these models is lower than for the models tested with the original dataset of Ref. [18]. This performance loss occurs due to the elimination of test-set data leakage, which biased the metrics upwards.^4^ This reduction highlights the importance of constructing test datasets in a manner that avoids the potential for information leakage to be able to produce accurate assessments of model performance [36] [37].

### 3.2 A Larger, Less Imbalanced Dataset Leads to Improved Models that Surpass Previously Published Models Even Further

After model training with the Ref. [18] datasets was complete, the dataset from Ref. [19] was located, which is more than an order of magnitude larger than the previously used dataset. Moreover, it contained a larger proportion of positive sites (2.44%), although the dataset was still significantly imbalanced.

Using a slightly modified training procedure (as described in Section 2.2), RNN models are trained on a modified version of the Ref. [19] dataset. Ref. [19] also included a potential motif for O-GlcNAcylation, which was tested on this modified dataset. Models with different window sizes were also tested to determine the effect of window sizes on predictive power. In the previous sections and in Ref. [18], window sizes were restricted to 10 AAs on each side of the S/T (21 AAs total), likely due to YinOYang’s fixed window size. However, it is reasonable to believe that AAs further away from the glycosylation site can have an effect on glycosylation; thus, models with up to 25 AAs on each side (51 AAs total) were investigated.

Our RNNs exhibit a much higher recall at the same precision level (Fig. 2) than any models in the previous sections (Figs. 1 and S1). Moreover, these RNNs have much higher precision for all recall values lower than 50%. As before, the precision for our RNN models increases monotonically with increasing threshold, while the F_1_ score peaks at different thresholds for each model. The performance of the models increases with increasing window size for sizes up to 20, and it also increases with increasing RNN hidden sizes for models with up to 225 neurons ^5^ (Fig. 2 and Table 3).

**Table 3:** Recall, precision, F_1_ score, and MCC metrics (in %) for the point with highest F_1_ score for models tested using a modified version of Ref. [19]’s dataset, and for the motif generated by Ref. [19]. Models “RNN-#” come from this work. The number that follows each RNN model is the LSTM module size used; the number after the semicolon represents the window size on each side of the central S/T. Model “YinOYang” is from Ref. [13]; its metrics are obtained by us. The “W-F Motifs” come from Ref. [19]. The number in parentheses represents the minimum number of amino acids (out of 5) that must follow the motif for a site to be considered positive. “No Model” sets all sites to potentially positive – that is, sets the threshold to 0.

| Metric | RNN-75; 5 win<br>(This Work) | RNN-75; 10 win<br>(This Work) | RNN-75; 15 win<br>(This Work) | RNN-75; 20 win<br>(This Work) |
| --- | --- | --- | --- | --- |
| Best Threshold | 0.90 | 0.95 | 0.95 | 0.80 |
| Recall (%) | 21.44 | 29.35 | 33.00 | 34.82 |
| Precision (%) | 16.20 | 24.22 | 29.38 | 29.87 |
| $F_1$ Score (%) | 18.45 | 26.54 | 31.09 | 32.16 |
| MCC (%) | 16.23 | 24.59 | 29.26 | 30.38 |

| Metric | RNN-150; 20 win<br>(This Work) | RNN-225; 20 win<br>(This Work) | YinOYang<br>(from [13]) | No Model |
| --- | --- | --- | --- | --- |
| Best Threshold | 0.50 | 0.60 | 0.60 | N/A |
| Recall (%) | 36.36 | 35.47 | 13.59 | 100 |
| Precision (%) | 34.58 | 36.90 | 8.08 | 2.44 |
| $F_1$ Score (%) | 35.45 | 36.17 | 10.13 | 4.76 |
| MCC (%) | 33.76 | 34.57 | 7.57 | 0.00 |

| Metric | W-F Motif (5)<br>(From [19]) | W-F Motif (4)<br>(From [19]) | W-F Motif (3)<br>(From [19]) | W-F Motif (2)<br>(From [19]) | W-F Motif (1)<br>(From [19]) |
| --- | --- | --- | --- | --- | --- |
| Recall (%) | 0.08 | 1.01 | 6.84 | 23.94 | 58.50 |
| Precision (%) | 39.29 | 20.33 | 11.85 | 5.89 | 3.38 |
| $F_1$ Score (%) | 0.16 | 1.92 | 8.67 | 9.45 | 6.38 |
| MCC (%) | 1.70 | 4.01 | 7.22 | 7.23 | 4.71 |

**Figure 2:**
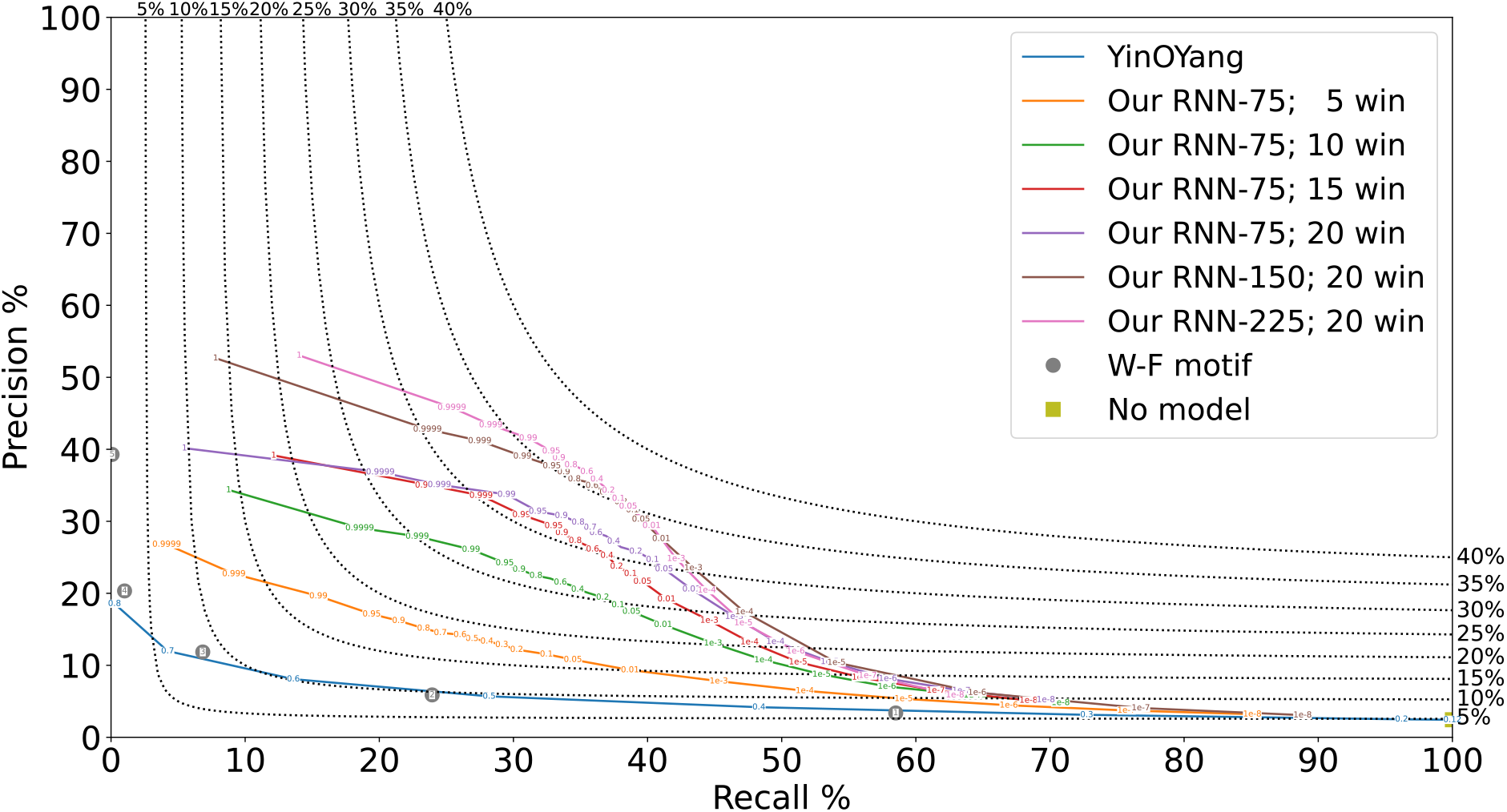
P-R curves for O-GlcNAcylation site prediction models tested using a modified version of Ref. [19]’s dataset. The black dotted lines are F_1_ score isolines, as labeled on the top and right sides of the figure. Model “YinOYang” is from Ref. [13]; its metrics are obtained by us. Models “Our RNN-#” come from this work. The number that follows each RNN model is the LSTM module size used; the number after the semicolon represents the window size on each side of the central S/T. The numbers on the “Our RNN-#” curves represent the minimum threshold for a site to be considered positive. The “W-F motif” comes from Ref. [19]; the numbers in the circles represent the minimum number of motif matches for a site to be considered positive. “No Model” sets all sites to potentially positive – that is, sets the threshold to 0.

Similarly to what occurred with the other datasets (Section 3.1 and Fig. S1), YinOYang performed very similarly to our models at low thresholds, and our models surpassed YoY at thresholds *≥* 10^*−*6^. Furthermore, YoY performed very similarly in this expanded dataset and in the smaller dataset from Section 3.1, indicating it is robust with respect to input data. However, its performance was also considerably low, surpassing an F_1_ score of 10% only at one threshold level. Even our RNN with a window size of only 5 AAs – that is, with half the information per sample – is Pareto dominant over YoY. With increasing window sizes, this disparity grows further, highlighting the quality of our RNNs and the positive impact from increasing window sizes. At its F_1_ score maximum, the RNN-225 model with a window size = 20 has a 357% improvement in precision, a 257% improvement in the F_1_ score, and a 357% improvement in MCC relative to YinOYang (Table 3).

To ensure our choice of cross-validation (later, training) and testing data is not biased, and to ensure the chosen hyperparameters are not excessively dependent on the particulars of a crossvalidation dataset, five-fold nested validation was performed on the RNN-225; 20 win model (as described in Section 2.1). The difference in performance among the five nested validation folds was negligible (Fig. S2 and Table S2), indicating that the chosen architecture and hyperparameters are robust to variations in the training and testing data. This highlights how this methodology is sound and can be applicable to any O-GlcNAcylation dataset. The best thresholds were also very similar for most folds; for the one fold where that was not the case, it should be noted that thresholds of 0.6 and 0.7 had the third- and second-best performance respectively, with absolute F_1_ score and MCC differences of less than 0.1%.

The previously proposed motif (from Ref. [19]) is excessively restrictive and fails to adequately capture the sequence needed for O-GlcNAcylation. Out of the 531,628 unique entries in the dataset, only 28 follow the motif, and only 11 out of those 28 are truly positive. Because this motif captured very few sequences overall, we hypothesized that allowing sequences to slightly deviate from the motif could lead to improved results. Allowing sequences to be treated as positive if at least 4 out of 5 amino acids follow the motif does not significantly improve the results: only 669 sites follow the motif, and only 136 out of those 669 are truly positive. The motif becomes slightly better when sequences are treated as positive if at least 3 out of 5 amino acids follow the motif; 7,787 sites follow the motif and 923 of those are true positives. If sequences are treated as positive if at least 2 out of 5 amino acids follow the motif, there are 54,875 positive sites and 3,232 true positives. Finally, treating sequences with just 1 out of 5 amino acids as positive leads to 233,943 positive sites and 7,898 true positives. While these deviations improved the motif’s results, the motif fails to achieve an F_1_ score or MCC greater than 10%, indicating it is not suitable to predict or describe O-GlcNAcylation (Table 3).

### 3.3 Interpretation of Model Predictions Using Shapley Values

The predictions of our models are interpreted via Shapley values by assigning a linear coefficient to each amino acid at each position based on a model’s predictions, leading to a (2 *×* window_size + 1) *×* 20 matrix of coefficients. Because each position has only one single amino acid, the final threshold for a given sequence is the sum of the (2 *×* window_size + 1) Shapley values of its amino acids. These allow for elucidation of the effect of each amino acid on the glycosylation chance (Fig. 3). The use of Shapley values with a threshold of 0.10–0.15 leads to minimal losses in performance, indicating the values are descriptive of the models’ predictions (Fig. S3 and Table S3).

**Figure 3:**
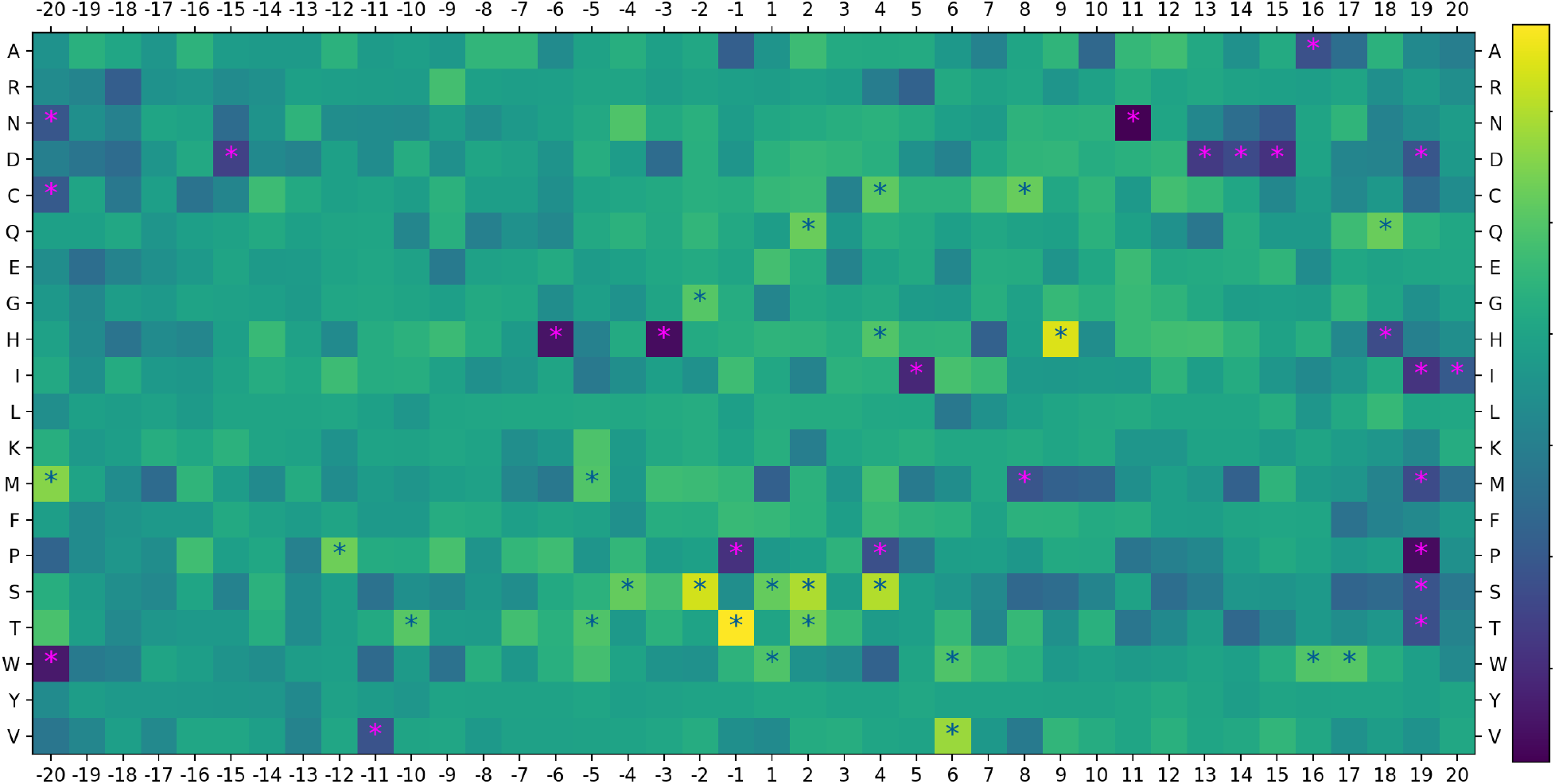
Heatmap of Shapley values for each amino acid and position from the RNN-225; 20 win model (Section 3.2). Yellow and light green squares indicate AA/position combinations that are more likely to be found in O-GlcNAcylated sequences; blue squares indicate AA/position combinations that are more likely to be found in non-O-GlcNAcylated sequences. Blue and magenta stars represents AA/position combinations at the top 3% or bottom 3% of values respectively. Units for this heatmap are arbitrary and thus not shown.

Some of the most relevant amino acid and position combinations are in agreement with Ref.[19]’s proposed motif and methodology (e.g.: a T at *−*1 or an A at 2). However, many others are in disagreement. For example, our RNN-225; 20 win considers an H at 9 or an S at *−*2 as considerably important, but these combinations are marginally less frequent in the positive samples relative to the negative samples. Some other amino acid and position combinations (such as a P at *−*1 or 4) are slightly more frequent in the positive samples relative to the negative samples, yet our model regards these combinations as critically negative for O-GlcNAcylation. These differences, combined with the superior performance of our models, indicate that the approach of simply counting the most frequent amino acids at each position (a unigram model) is not adequate to describe and predict O-GlcNAcylation, and it is likely that combinatorial effects (such as the secondary structure of, or the net charge near the potential site) play an important role in O-GlcNAcylation.

## 4 Discussion

This work constructs MLP and RNN models from multiple sources of literature data on protein O-GlcNAcylation based on human-selected descriptors (first part of Section 3.1) or protein sequences (second part of Section 3.1 and all of Section 3.2). An MLP model trained in this work was compared with previously published models (as reported in Ref. [18]), and this work’s MLP model surpasses the previously published models in precision (307% improvement), F_1_ score (151% improvement), and MCC (90.9% improvement) metrics (Section 3.1). This study contrasts with the past studies by our use of different model architectures, multiple prediction thresholds, and rigorous cross-validation for hyperparameter selection. An analysis of the dataset of Ref. [18] found multiple issues, however, including test-set leakage, making the results overoptimistic for all models.

To address the dataset issues, a new dataset is constructed from the original dataset of Ref. [18]. This new dataset contains protein sequences instead of descriptors, promoting the training of RNN models and simplifying overall usage for the end-users. Two RNN models trained in this work are compared with YinOYang [13], the only model evaluated in Ref. [18] that is still available (second part of Section 3.1). The correction of the dataset’s issues lowered the overall performance of the models tested. Nevertheless, the best RNN model still displayed a 391% improvement in precision, 134% improvement in F_1_ score, and 136% improvement in MCC when compared to YinOYang, and the RNN models surpassed an MLP model trained on the same data (Fig. S1). This comparison suggests that the RNN model architecture is better suited for prediction of O-GlcNAcylation than the MLP model architecture, which is consistent with the RNN being able to leverage the sequential data structure of protein site sequences instead of the MLP’s limited way of handling structured data.

A much larger dataset from Ref. [19] is refined and used to train RNN models through a slightly modified methodology. These RNN models displayed superior performance when compared to the other models in previous works, displaying a 357% improvement in precision, 257% improvement in F_1_ score, and 357% improvement in MCC over YinOYang (Section 3.2). These RNN models were also better than the RNN models trained in the previous section, and we hypothesize part of this difference is due to the greater number of entries and lower data imbalance found in Ref. [19]’s modified dataset. The use of weight decay and learning rate scheduling contributed to further improving the performance of the RNN models in Section 3.2.

Ref. [19] also proposes a motif for O-GlcNAcylation, but that motif is excessively restrictive. The original formulation of Ref. [19] motif has an F_1_ score of only 0.16% and an MCC of 1.70%, indicating it is barely better than random guessing. While making it less strict increased its performance, that motif never achieves an F_1_ score or MCC *≥* 10%. The Shapley values extracted from our models, on the other hand, provide interpretability while maintaining most of the superior performance of our models (Section 3.3, Fig. S3, and Table S3). While a few of the most positive or negative Shapley values match the motif proposed in Ref. [19], many others do not. Given the higher performance of the Shapley value predictions over Ref. [19]’s motif, this suggests a simple unigram model is not adequate to describe and predict O-GlcNAcylation, and more complex models that take into account interactions are necessary.

This software used in this work is publicly available, allowing other researchers to reproduce this work and reuse or improve the code in future studies. The software provides a simple way to install and run the best RNN model (trained on Ref. [19]’s modified data and using a window size of 20 AAs on each side of the central S/T) to predict the presence of O-GlcNAcylation sites based on the local protein sequence. Instructions are provided in Sections S1 and S2 of the supplemental data or in the README in our GitHub repository.

## Data availability statement

The raw data, code used to process these data, and models underlying this article are available in a GitHub repository at https://github.com/PedroSeber/O-GlcNAcylation_Prediction

## Author Contributions

**Pedro Seber**: methodology, data curation, software, validation, formal analysis, writing – original draft, writing – review & editing, visualization.

**Richard D. Braatz**: conceptualization, methodology, resources, writing – original draft, writing – review & editing, supervision, project administration, funding acquisition.

## Supplemental Information

### S1 Reproducing the models and plots

The models can be recreated by downloading the datasets and running the `ANN_train.py` file with the appropriate flags (run `python ANN_train.py --help` for details).

The plots can be recreated by running the `make_plot.py` file with the appropriate data version as an input (`python make_plot.py v1` for Fig. 1, `python make_plot.py v5` for Fig. 2, `python make_plot.py shap_heatmap` for Fig. 3, `python make_plot.py v3` for Fig. S1, `python make_plot.py v5_nested` for Fig. S2, and `python make_plot.py v5_shap` for Fig. S3).

### S2 Using the RNN model to predict O-GlcNAcylation sites

The Conda environment defining the specific packages and version numbers used in this work is available as `ANN_environment.yaml` on our GitHub (github.com/PedroSeber/O-GlcNAcylation_Prediction). To use our trained model, run the `Predict.py` file as `python Predict.py <sequence> -t <threshold> -bs <batch_size>`.

Alternatively, create an (N+1)x1 .csv with the first row as a header (such as “Sequences”) and all other N rows as the actual amino acid sequences, then run the Predict.py file as `python ANN_predict.py <path/to/file.csv> -t <threshold> -bs <batch_size>`. Results are saved as a new .csv file.

To run Shapley value predictions in addition to the model predictions, run the `Predict.py` file with the `-shap` flag. Whatever other flags should be included as needed.

### S3 Equations for the model evaluation metrics (Section 2.3)

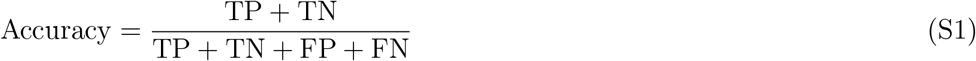

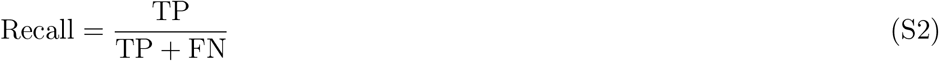

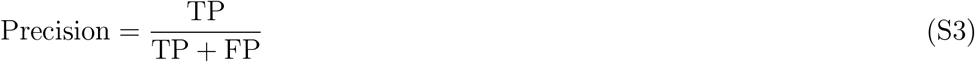

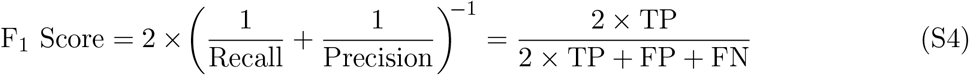

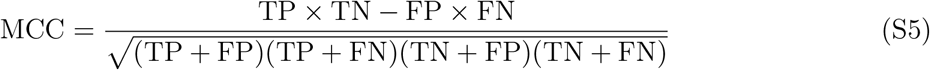

### S4 Supplemental Figures & Tables

**Table S1:** Recall, precision, F_1_ score, and MCC metrics (in %) for the points with highest F_1_ score for O-GlcNAcylation site prediction models tested on the modified dataset of Ref. [18]. Model “YinOYang” is from Ref. [13]; its metrics are obtained by us. As mentioned above, other previously published models are not available; thus, their performance on this dataset could not be obtained. Models “Our RNN” come from this work. The number that follows each of our RNN models is the LSTM module size. “No Model” sets all sites to potentially positive – that is, sets the threshold to 0.

| Metric | YinOYang<br>(From [13]) | Our RNN-76<br>(This Work) | Our RNN-152<br>(This Work) | No Model |
| --- | --- | --- | --- | --- |
| Recall (%) | 21.50 | 15.31 | 13.27 | 100 |
| Precision (%) | 4.93 | 24.19 | 24.53 | 1.29 |
| $F_1$ Score (%) | 8.02 | 18.75 | 17.22 | 2.55 |
| MCC (%) | 7.83 | 18.48 | 16.83 | 0.00 |

**Figure S1:**
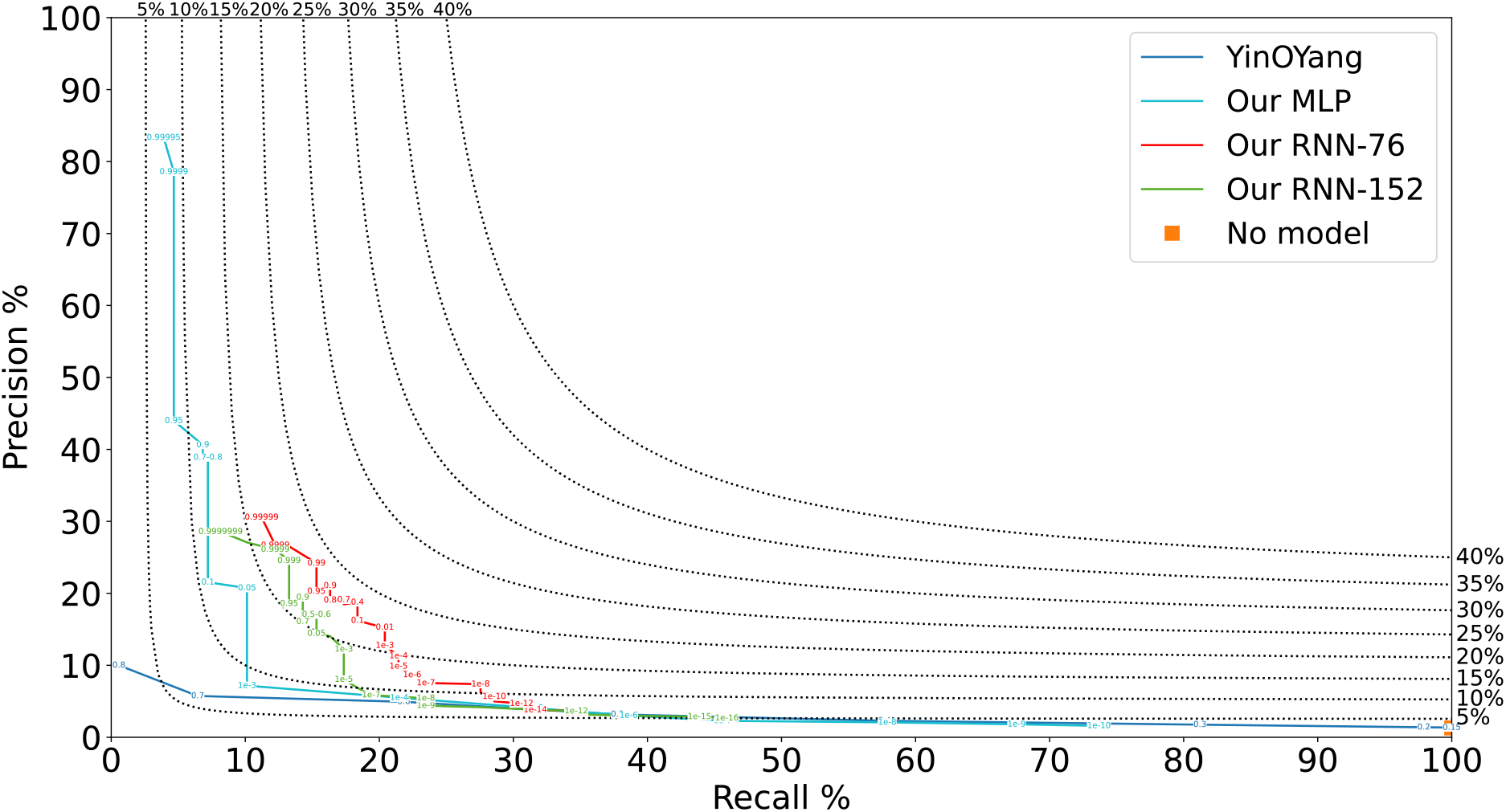
Precision-recall curves for O-GlcNAcylation site prediction models tested on the modified dataset of Ref. [18]. The black dotted lines are F_1_ score isolines, as labeled on the top and right sides of the figure. Model “YinOYang” is from Ref. [13]; its metrics are obtained by us. As mentioned in Section 3.1, other previously published models are not available; thus, their performance on this dataset could not be obtained. Models “Our MLP” and “Our RNN” come from this work. The number that follows each of our RNN models is the LSTM module size used. The numbers on each curve represent the minimum threshold for a site to be considered positive. “No Model” sets all sites to potentially positive – that is, sets the threshold to 0.

**Table S2:** Recall, precision, F_1_ score, and MCC metrics (in %) for the point with highest F_1_ score for the RNN-225; 20 win model after nested validation on a modified version of Ref. [19]’s dataset

| Metric | Fold 1 | Fold 2 | Fold 3 | Fold 4 | Fold 5 | Mean ($\pm\sigma$) |
| --- | --- | --- | --- | --- | --- | --- |
| Best Threshold | 0.60 | 0.70 | 0.70 | 0.60 | 0.95 | N/A |
| Recall (%) | 35.47 | 36.08 | 35.00 | 36.48 | 33.46 | $35.30\pm1.05$ |
| Precision (%) | 36.90 | 34.27 | 36.33 | 33.76 | 38.01 | $35.85\pm1.60$ |
| $F_1$ Score (%) | 36.17 | 35.16 | 35.65 | 35.07 | 35.59 | $35.53\pm0.39$ |
| MCC (%) | 34.57 | 33.52 | 34.05 | 33.45 | 34.15 | $33.95\pm0.42$ |

**Figure S2:**
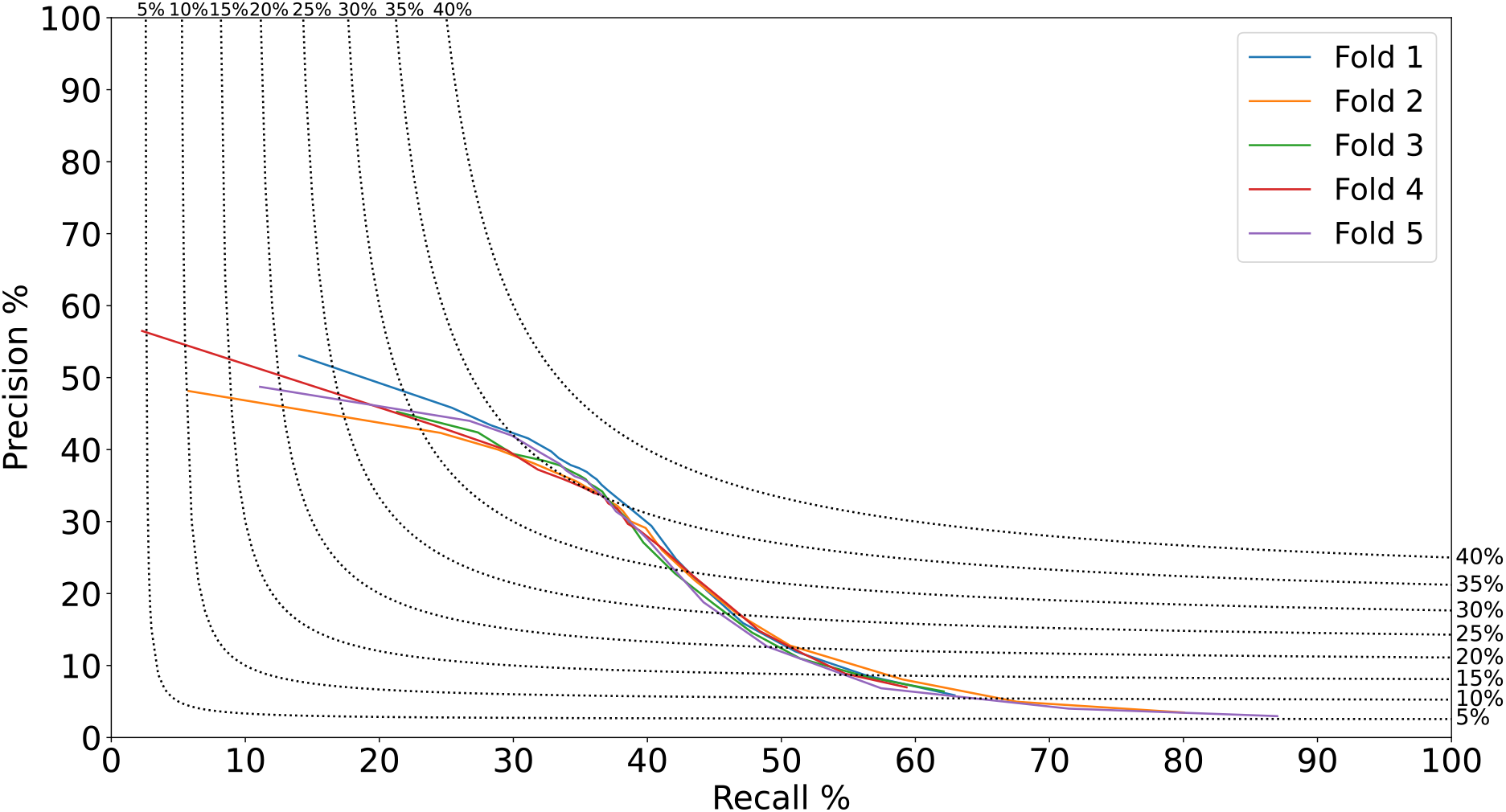
Precision-recall curves for the RNN-225; 20 win model after nested validation on a modified version of Ref. [19]’s dataset. The black dotted lines are F_1_ score isolines, as labeled on the top and right sides of the figure.

**Table S3:**
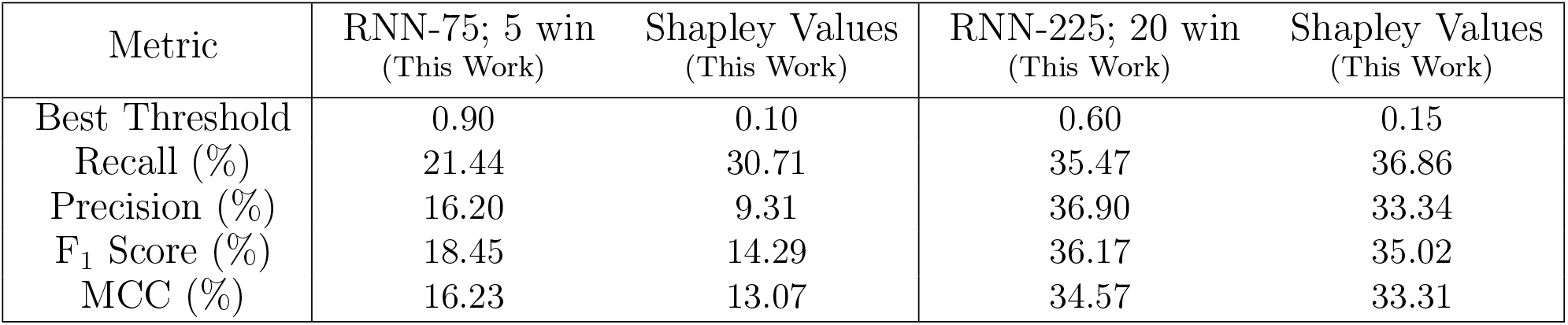
Recall, precision, F_1_ score, and MCC metrics (in %) for the point with highest F_1_ score for models tested using a modified version of Ref. [19]’s dataset. Models “RNN-#” come from this work. The number that follows each RNN model is the LSTM module size used; the number after the semicolon represents the window size on each side of the central S/T (see also Table 3). Immediately after are the predictions from Shapley values from these RNN models.

| Metric | RNN-75; 5 win<br>(This Work) | Shapley Values<br>(This Work) | RNN-225; 20 win<br>(This Work) | Shapley Values<br>(This Work) |
| --- | --- | --- | --- | --- |
| Best Threshold | 0.90 | 0.10 | 0.60 | 0.15 |
| Recall (%) | 21.44 | 30.71 | 35.47 | 36.86 |
| Precision (%) | 16.20 | 9.31 | 36.90 | 33.34 |
| $F_1$ Score (%) | 18.45 | 14.29 | 36.17 | 35.02 |
| MCC (%) | 16.23 | 13.07 | 34.57 | 33.31 |

**Figure S3:**
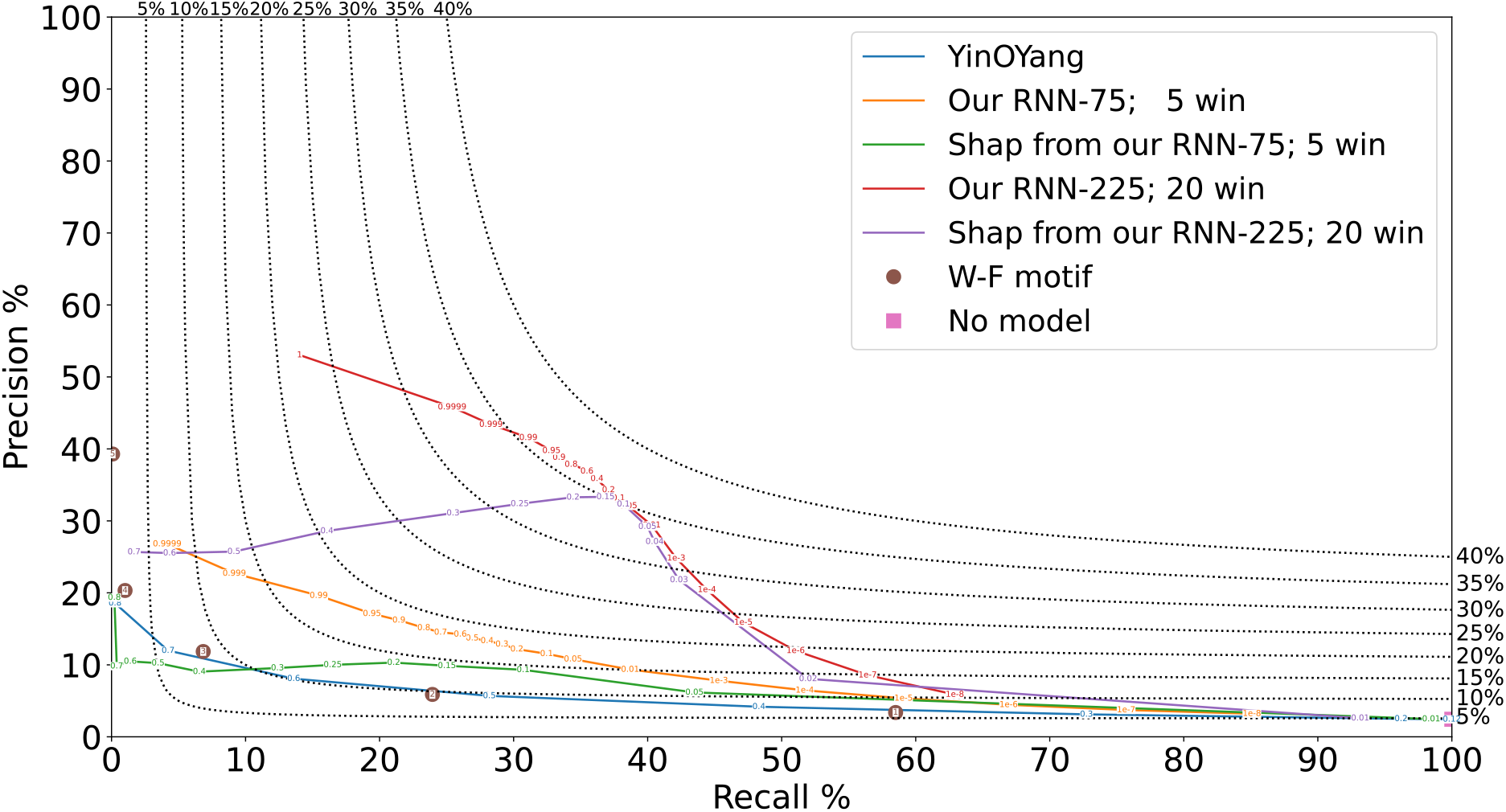
P-R curves for O-GlcNAcylation site prediction models tested using a modified version of Ref. [19]’s dataset. The black dotted lines are F_1_ score isolines, as labeled on the top and right sides of the figure. Model “YinOYang” is from Ref. [13]; its metrics are obtained by us. Models “Our RNN-#” come from this work. The number that follows each RNN model is the LSTM module size used; the number after the semicolon represents the window size on each side of the central S/T. Immediately below are the predictions from Shapley values from these RNN models. The numbers on the “Our RNN-#” curves represent the minimum threshold for a site to be considered positive. The “W-F motif” comes from Ref. [19]; the numbers in the circles represent the minimum number of motif matches for a site to be considered positive. “No Model” sets all sites to potentially positive – that is, sets the threshold to 0.

## Footnotes

1 As detailed in the `ANN_train.py` file in the GitHub repository.

2 RNNs with 38 and 228 neurons had cross-validation results inferior to RNN-76 and RNN-152, and so are not included.

3 O-GlcNAcPRED-II and OGTSite are no longer available; thus, their metrics on the Modified dataset cannot be obtained.

4 The performance loss was not due to feature changes. MLPs using this dataset and the same features lost more than 10% of their absolute recall values at the same precision and performed worse than the RNNs (Fig. S1).

5 Models with window sizes = 25 or RNN sizes = 300 neurons performed nearly identically to models with window sizes = 20 or RNN sizes = 150 or 225 neurons, so the former are not included in Fig. 2 or Table 3.

